# Segregation between the parietal memory network and the default mode network: Effects of spatial smoothing and model order in ICA

**DOI:** 10.1101/086454

**Authors:** Yang Hu, Jijun Wang, Chunbo Li, Yin-shan Wang, Zhi Yang, Xi-Nian Zuo

## Abstract

A brain network consisting of two key parietal nodes, the precuneus and the posterior cingulate cortex, has emerged from recent fMRI studies. Though it is anatomically adjacent to and spatially overlaps with the default mode network (DMN), its function has been associated with memory processing, and it has been referred to as the parietal memory network (PMN). Independent component analysis (ICA) is the most common data-driven method of extracting PMN and DMN simultaneously. However, the effects of data preprocessing and parameter determination in ICA on PMN-DMN segregation are completely unknown. Here, we employ three typical algorithms of group ICA to assess how spatial smoothing and model order influence the degree of PMN-DMN segregation. Our findings indicate that PMN and DMN can only be stably separated using a combination of low-level spatial smoothing and high-order model across the three ICA algorithms. We thus argue for more considerations on parametric settings for interpreting DMN data.

## Introduction

Resting state networks (RSN) refer to a set of brain regions acting in a similar fashion without a specific task or stimulus, of which, the default mode network (DMN) is the most heavily investigated RSN [1, 2]. Recently, an RSN anchored at two parietal areas including the precuneus and the posterior cingulate has been identified in various studies of DMN. Despite their spatial proximity, the separation of the posterior parietal network from DMN is important from a neuroscience perspective. Specifically, this network has a different developmental trajectory from DMN [3] and is named the parietal memory network (PMN) due to its functional role in novel memory-related processing [4]. A comprehensive review of the functional anatomy of DMN white matter connections and neuropsychological findings ruled-out the precuneus as a structure within the DMN [1]. The segregation between PMN and DMN has also been revealed in various studies examining low-frequency fluctuations in spontaneous brain activity as measured by resting-state functional MRI (rfMRI) [5-7]. In practice, however, PMN can be easily mislabeled as a posterior part of DMN. Due to the spatial proximity and various methodological issues, the interpretation of DMN-related neuroimaging findings is difficult.

Several methods including clustering, graph theory and independent component analysis (ICA) have been used to extract PMN and DMN [5, 7, 8]. ICA can obtain multiple spatially overlapped RSNs without a priori knowledge and is therefore perfectly suitable for examining PMN/DMN activities, as the boundaries between these two networks are not well-defined. Previous studies have demonstrated that model order (MO) can significantly impact the estimated RSNs while a high MO was recommended for ICA [9, 10]. Of note, these studies did not directly evaluate the effects of MO on PMN-DMN segregation. In addition, spatial smoothing (SM) is a common step in rfMRI preprocessing prior to group ICA analysis [11, 12]. SM increases the signal-to-noise ratio and accounts for inter-individual registration bias. However, SM with a larger Gaussian kernel could decrease the spatial resolution and potentially blur the signals from functionally distinct areas [13, 14]. Because the PMN surrounds the posterior part of the DMN, the influence of SM kernel size should not be overlooked. Unfortunately, to the best of our knowledge, the effects of MO and SM on PMN-DMN segregation have not been systematically investigated.

Currently, there are three different types of ICA algorithms used in group-level rfMRI data analysis: (1) apply ICA to each individual dataset without any constraints on the dependence among individuals and combine individual results with different similarity metrics post-hoc [3, 15, 16]; (2) apply ICA to each individual dataset and simultaneously take the inter-individual dependence into consideration [17, 18]; (3) transform all individual-level datasets into one group-level dataset, and then apply ICA to the aggregated dataset with an additional procedure of back-reconstruction of individual components [19-22]. The answer to the question of whether PMN-DMN segregation using ICA is algorithm-dependent is not trivial.

To determine the optimal parametric settings for group ICA for robust PMN-DMN separation, in this study, we aim to systematically investigate the effects of MO and SM on the segregation of these two networks using three types of group level ICA methods. Using various MO-SM combinations in these three algorithms, we identified components representing PMN and DMN using spatial templates and a series of objective criteria. We evaluated the quality of the PMN and DMN identified by ICA using goodness-of-fit, mean weights, and inter-individual reproducibility. We hypothesized that both MO and SM would affect PMN-DMN discrimination.

## Materials and Methods

### Subjects

Sixty-five healthy subjects (age 25.13 ± 6.42 years, 30♂, 35♀) were recruited from Shanghai Mental Health Center. The Institutional Review Board at the Shanghai Mental Health Center approved the study protocol. Written informed consent was obtained from each participant or the participant’s guardian prior to data acquisition. The inclusion criteria for the healthy subjects were as follows: (1) age ranging from 15-40; (2) no serious physical diseases, pregnancy, or substance abuse; (3) no psychoactive substance use for at least one month; (4) no history of mental disorder; (5) education levels exceeding primary school level. The exclusion criteria for healthy controls were as follows: (1) meet the criteria for any mental disorder according to DSM-IV; (2) family history of mental disorder; (3) unstable mental state; (4) history of taking antipsychotic drugs; (5) substance abuse in the past month; (6) pregnancy; (7) history of serious physical disease; (8) unsuitability for MRI scans.

### Data acquisition

All imaging data were collected using a 3.0 Tesla Siemens Verio MRI scanner (Enlargen, Germany) at the Shanghai Mental Health Center. Resting-state scans were acquired with an echo-planar imaging (EPI) sequence (45 axial interleaved slices, acquired from inferior to superior, FOV = 216 mm, matrix = 72x72, slice thickness/gap = 3.0/0.0 mm, no gap, TR/TE = 3000/30 ms, flip angle = 85°, 170 volumes, duration 8’30’’). High-resolution anatomical scans were acquired with a T1-weighted 3D MP-RAGE sequence (192 sagittal slices, FOV = 256 mm, matrix = 256x240, slice thickness/gap = 1.0/0.0 mm, TR/TE/TI = 2300/2.96/900 ms). Participants were instructed to close their eyes and remain awake during scanning.

### Preprocessing and quality control

Both anatomical and rfMRI images were preprocessed using the Connectome Computation System [23]. Structural images were first cleaned by using a spatially adaptive non-local mean filter to remove noise [24] and fed into FreeSurfer [25] for extracting the brain as well as for segmenting the brain tissues into gray matter, white matter and cerebrospinal fluid. All images were converted into MNI152 space using Advanced Normalization Tools (ANTs) [26].

The following preprocessing steps were applied to the rfMRI images: (1) the first 15 volumes were discarded to allow MRI signal equilibration; (2) slice timing differences were corrected; (3) the head movements were realigned over the entire scan; (4) the mean rfMRI image was spatially normalized to MNI152 space via the combined registration of a rigid transformation of the individual structural images and nonlinear ANTs transformation; (5) the 4D data were standardized to a global mean intensity of 10,000; (6) the data were temporally band-pass (0.01-0.1Hz) filtered; (7) the data were then spatially smoothed using a 0, 6, 9 and 12 mm FWHM Gaussian kernel.

The quality of the brain extraction and registration were visually inspected. The anatomical images of four participants were excluded from further analysis due to poor brain extraction or registration quality. Head motion in the rfMRI data of the 61 participants was evaluated using mean frame-wise displacement (meanFD)[27], and the meanFD was less than 0.2 mm.

### Group ICA analysis

We chose Generalized Ranking and Averaging ICA by Reproducibility (gRAICAR) [3, 28], Independent Vector Analysis (IVA-GL) [17, 18] and Time-Concatenated Group ICA (TCgICA) [21], corresponding to the three types of group ICA mentioned above, to perform brain network extraction from the preprocessed rfMRI datasets. The principle behind each algorithm will be generally introduced, followed by a detailed description of the parameter settings and the analysis steps.

gRAICAR (https://github.com/yangzhi-psy/gRAICAR) is a matching algorithm performed on a group of independent components (ICs) derived from the individual-level ICA based on similarity. Each preprocessed rfMRI dataset was decomposed into a number of ICs in individual native space using MELODIC [29]. The ICs were then transformed into MNI152 standard space. The individual-level and normalized ICs were further fed into the gRAICAR algorithm for alignment across participants. Finally, a weighted average of the aligned ICs was computed and then Z-transformed (zero mean and one standard deviation) to produce representative group-level ICs.

IVA-GL decomposes all of the individual-level datasets simultaneously by assuming a multivariate probability density function to maximize inter-individual linear and non-linear dependence. IVA-GL was implemented using the GIFT toolbox (http://mialab.mrn.org/software/gift). Specifically, the individual preprocessed rfMRI datasets were temporally de-meaned then processed with IVA-GL to decompose them into individual-level ICs all at once. The z-scores of the individual-level ICs were calculated, averaged and Z-transformed to obtain the group-level ICs.

Temporal concatenation group ICA (TCgICA) assumes all participants share a set of common RSNs to be extracted. The processing steps of TCgICA include: (1) all individual rfMRI datasets were temporally concatenated into a large 4D file, which was then reduced using principle component analysis to obtain a group-level dataset; (2) the group dataset was then decomposed into a set of group-level ICs; (3) individual-level ICs were finally back-reconstructed using dual-regression [20, 30]. TCgICA and dual-regression were implemented and carried out in MELODIC (fsl.fmrib.ox.ac.uk/fsl/fslwiki/MELODIC).

The above analyses were conducted on preprocessed rfMRI datasets using four different SM kernel sizes (0, 6, 9, 12 mm). Five different MOs were specified (20, 40, 60, 80, 100), yielding 20 (SM, MO) combinations or sets of group-level ICs and their corresponding individual-level ICs for each algorithm. Of note, in assessing the effects of the parameter settings on PMN-DMN segregation, the MO settings for the individual-level or group-level analyses depended on the algorithm used.

### PMN/DMN selection

PMN and DMN were automatically selected using a template-matching scheme. Spatial templates for PMN and DMN were generated based on a 17-network parcellation of human cerebral cortex [7]. Regarding the observation that DMN tended to split into anterior and posterior sub-networks in the high-MO condition [10], we manually sectioned the DMN template into anterior DMN (aDMN) and posterior DMN (pDMN). The pDMN has three clusters located in the precuneus, posterior cingulate and bilateral angular gyrus. To better characterize the anatomical specificity of PMN and DMN, we set anchor points in the center of the mid-line clusters in PMN and pDMN. Figure 1 illustrates the spatial templates of both PMN and DMN with the anchor points.

**Fig. 1.**
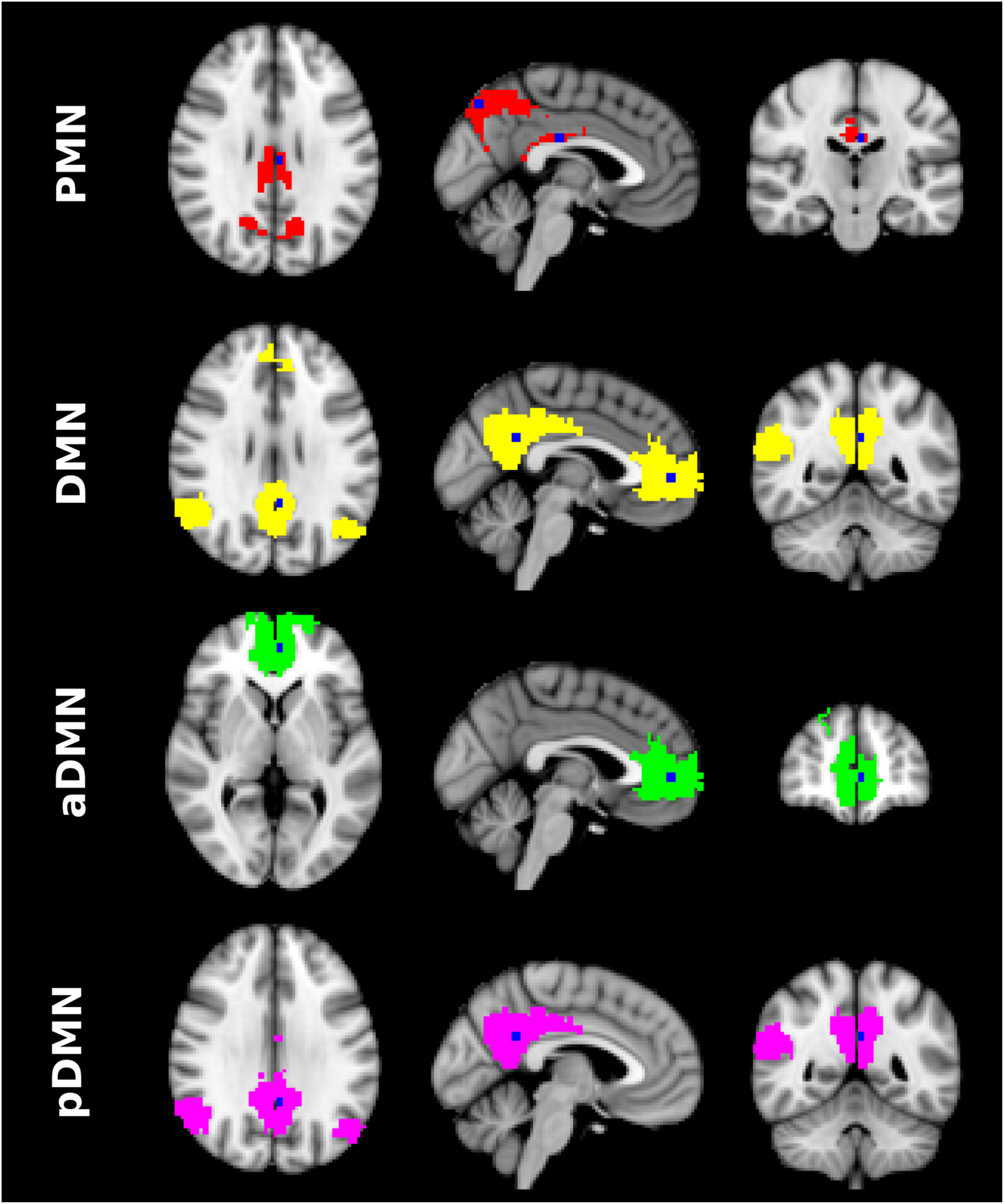
PMN and DMN templates with corresponding anchor points. PMN template in red and anchor points in blue, located in precuneus and posterior cingulate. DMN template in yellow and anchor points in blue, located in posterior cingulate and paracingulate gyrus. Anterior DMN template in green and anchor point in blue, located in paracingulate gyrus. Posterior DMN template in pink and anchor point in blue, located in posterior cingulate.

For each (SM, MO) combination, the following IC selection procedure was carried out (see Figure 2 for a diagram on the selection procedure): (1) three ICs were selected from the set of group-level ICs that were most highly correlated with the PMN, DMN and pDMN templates; (2) the candidate ICs were then thresholded at Z > 2 for gRAICAR and IVA-GL, and at Z > 5 for TCgICA; (3) the anchor points were assessed to determine if they were included in the thresholded maps of the ICs representing PMN/DMN/pDMN. If the candidate ICs of PMN failed to include the anchor points, PMN was labeled as “Not Found”. If the candidate ICs of DMN and pDMN failed to include the corresponding anchor points, DMN was labeled as “Not Found”; (4) the ICs representing PMN and DMN/pDMN were assessed to determine if they are the same IC. If so, the IC was assigned to the RSN with the higher correlation coefficient and the other was labeled “Not Found”.

**Fig. 2.**
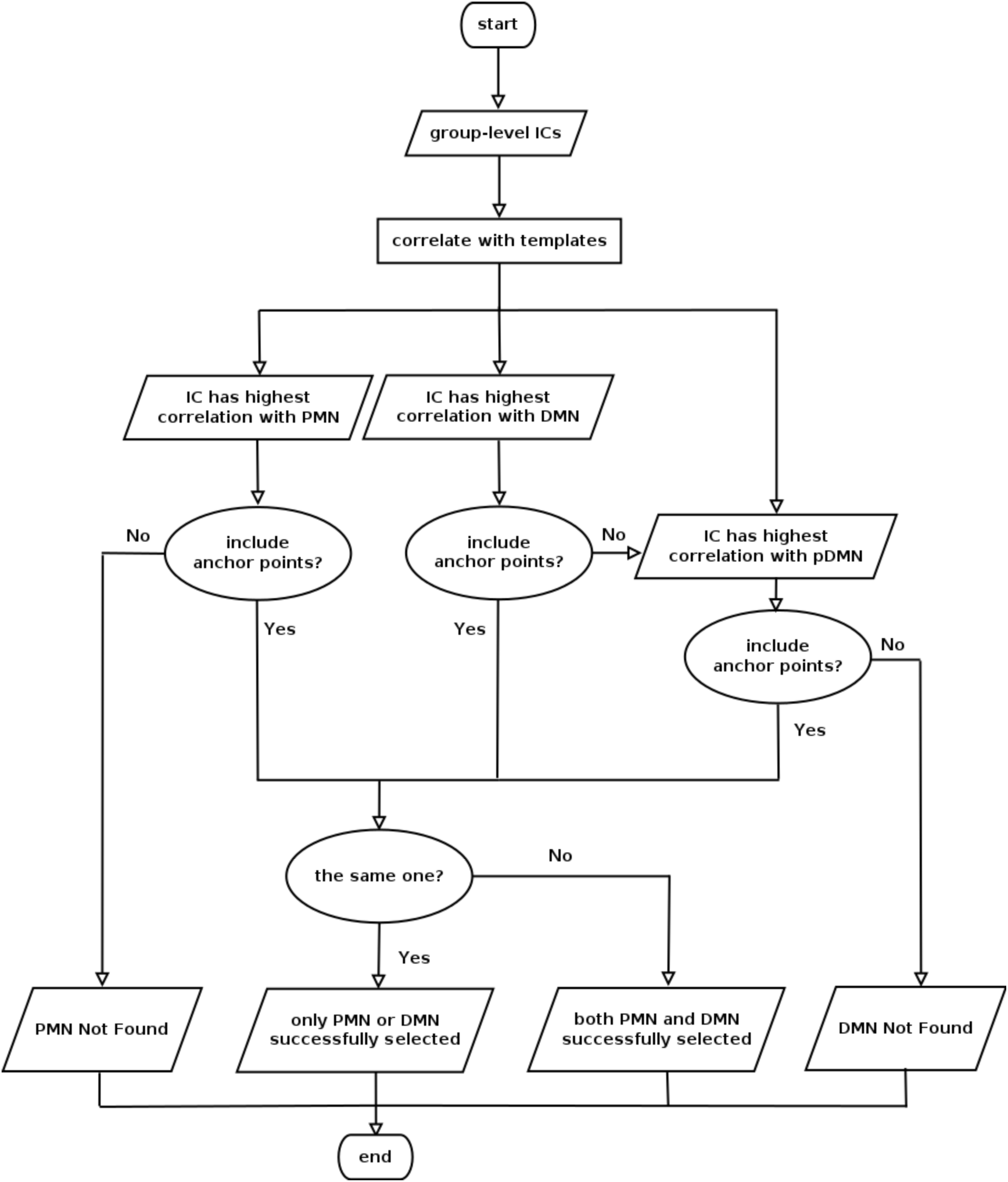
Flow chart presenting procedures for selecting independent components (ICs) corresponding to PMN and DMN.

In addition to the selection criteria described above, Supplementary Figure 1-3 present the ICs simply selected by the highest correlation with PMN, DMN, aDMN, and pDMN templates to provide a more complete picture of the selected ICs. Assessing the ICs in this way, we can determine whether PMN and posterior DMN were truly independent from each other.

### Goodness-of-fit, mean weights, and inter-individual similarity

Goodness-of-fit (GoF) was defined as the Pearson correlation coefficient between the selected IC and the corresponding template. A higher goodness-of-fit indicates better correspondence between the template and the selected IC. We further characterized the selected ICs representing PMN and DMN by measuring their mean weights and inter-individual similarity.

Mean weights (MW) were calculated by thresholding the selected ICs and averaging the weights of the remaining voxels. A higher mean weight reflects a clearly defined RSN. Conversely, a low mean weight indicates that the RSN was not specifically represented by the selected IC. Inter-individual similarity (IIS) was defined as the mean and standard deviation of the Pearson correlation coefficients from the un-thresholded spatial maps of the individual-level ICs. High mean inter-individual similarity indicates that the represented RSNs were consistent across participants.

## Results

The selected ICs representing PMN and DMN, the GoF scores, MW and IIS are presented in Tables 1-3, corresponding to the results from gRAICAR, IVA-GL, and TCgICA. To summarize the results, we grouped our results into four conditions of SM/MO combinations: (low SM 0/6, low MO 20/40), (low SM 0/6, high MO 80/100), (high SM 9/12, low MO 20/40) and (high SM 9/12, high MO 80/100). Figure 3 presents the four most extreme cases, i.e., SM 0/MO 20, SM 0/MO 100, SM 12/MO 20, and SM12/MO 100 as examples. For a full description of all 20 conditions across the three algorithms, please refer to the Supplementary Figures 1-3.

**Fig. 3.**
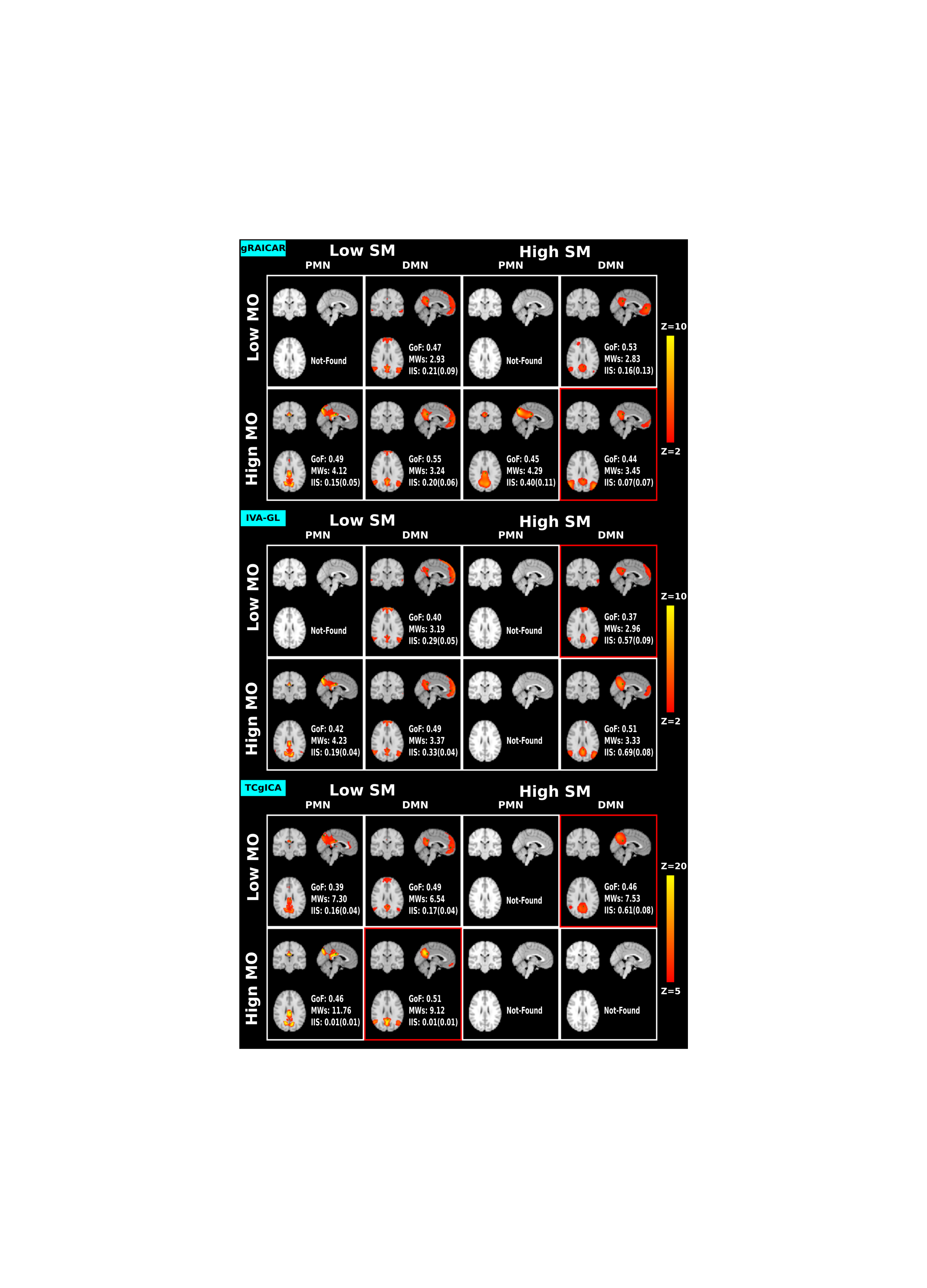
The selected IC maps representing PMN and DMN in four combinations of spatial smoothing levels and model orders across three algorithms. A red border means that the RSN is identified as posterior DMN.

**Table 1.** The GoF, MWs and IIS of selected PMN/DMN under different combinations of model order and spatial smoothing levels using gRAICAR.

| gRAICAR |  | SM 0 |  | SM 6 |  | SM 9 |  | SM 12 |  |
| --- | --- | --- | --- | --- | --- | --- | --- | --- | --- |
|  |  | PMN | DMN | PMN | DMN | PMN | DMN | PMN | DMN |
| MO 20 | GoF | – | 0.47 | – | 0.51 | – | 0.47 | – | 0.53 |
|  | MWs | – | 2.92 | – | 2.87 | – | 2.94 | – | 2.82 |
|  | IIS<sup>a</sup> | – | 0.21(0.09) | – | 0.30(0.12) | – | 0.30(0.14) | – | 0.16(0.12) |
| MO 40 | GoF | 0.40 | 0.55 | 0.48 | 0.52 | 0.47 | – | 0.45 | 0.49 |
|  | MWs | 3.42 | 3.18 | 3.83 | 3.16 | 3.81 | – | 3.80 | 3.52 |
|  | IIS<sup>a</sup> | 0.05(0.05) | 0.20(0.08) | 0.28(0.11) | 0.28(0.11) | 0.36(0.13) | – | 0.41(0.13) | 0.35(0.14) |
| MO 60 | GoF | 0.47 | 0.56 | 0.50 | 0.57 | 0.49 | 0.57 | 0.47 | – |
|  | MWs | 3.91 | 3.26 | 4.09 | 3.30 | 4.08 | 3.19 | 4.13 | – |
|  | IIS<sup>a</sup> | 0.10(0.07) | 0.22(0.07) | 0.33(0.08) | 0.28(0.10) | 0.39(0.11) | 0.22(0.13) | 0.43(0.13) | – |
| MO 80 | GoF | 0.49 | 0.56 | 0.50 | 0.55 | 0.47 | 0.43 | 0.45 | – |
|  | MWs | 4.14 | 3.27 | 4.22 | 3.28 | 4.27 | 3.81 | 4.29 | – |
|  | IIS<sup>a</sup> | 0.15(0.06) | 0.20(0.07) | 0.30(0.07) | 0.22(0.09) | 0.25(0.08) | 0.06(0.05) | 0.18(0.09) | – |
| MO 100 | GoF | 0.49 | 0.55 | 0.49 | 0.54 | 0.48 | 0.51 | 0.45 | 0.44 |
|  | MWs | 4.12 | 3.23 | 4.21 | 3.31 | 4.26 | 3.37 | 4.29 | 3.45 |
|  | IIS<sup>a</sup> | 0.14(0.05) | 0.20(0.06) | 0.31(0.07) | 0.21(0.09) | 0.38(0.08) | 0.08(0.08) | 0.40(0.10) | 0.06(0.06) |

<sup>a</sup>Data presented as: mean (standard deviation);
“–” indicates the corresponding IC was not found.

**Table 2.** The GoF, MWs and IIS of selected PMN/DMN under different combinations of model order and spatial smoothing levels using IVA-GL.

| IVA-GL |  | SM 0 |  | SM 6 |  | SM 9 |  | SM 12 |  |
| --- | --- | --- | --- | --- | --- | --- | --- | --- | --- |
|  |  | PMN | DMN | PMN | DMN | PMN | DMN | PMN | DMN |
| MO 20 | GoF | – | 0.40 | – | 0.36 | – | 0.36 | – | 0.37 |
|  | MW<sub>s</sub> | – | 3.19 | – | 3.07 | – | 3.02 | – | 2.96 |
|  | IIS<sup>a</sup> | – | 0.29(0.05) | – | 0.49(0.05) | – | 0.52(0.08) | – | 0.57(0.09) |
| MO 40 | GoF | – | 0.45 | 0.46 | – | 0.46 | 0.35 | – | 0.37 |
|  | MW<sub>s</sub> | – | 3.22 | 3.90 | – | 3.99 | 3.12 | – | 3.55 |
|  | IIS<sup>a</sup> | – | 0.29(0.05) | 0.47(0.06) | – | 0.43(0.09) | 0.52(0.07) | – | 0.44(0.10) |
| MO 60 | GoF | 0.46 | 0.33 | 0.52 | 0.38 | 0.52 | – | 0.50 | 0.37 |
|  | MW<sub>s</sub> | 4.01 | 3.16 | 4.21 | 3.16 | 4.27 | – | 4.26 | 3.43 |
|  | IIS<sup>a</sup> | 0.27(0.04) | 0.31(0.04) | 0.43(0.06) | 0.45(0.06) | 0.55(0.06) | – | 0.57(0.08) | 0.61(0.06) |
| MO 80 | GoF | 0.45 | 0.35 | 0.52 | 0.50 | 0.51 | – | 0.48 | – |
|  | MW<sub>s</sub> | 4.27 | 3.23 | 4.37 | 3.44 | 4.44 | – | 4.49 | – |
|  | IIS<sup>a</sup> | 0.19(0.04) | 0.30(0.04) | 0.45(0.06) | 0.44(0.06) | 0.58(0.06) | – | 0.66(0.05) | – |
| MO 100 | GoF | 0.42 | 0.49 | 0.51 | 0.56 | 0.50 | 0.37 | – | 0.51 |
|  | MW<sub>s</sub> | 4.23 | 3.37 | 4.26 | 3.36 | 4.30 | 4.30 | – | 3.33 |
|  | IIS<sup>a</sup> | 0.19(0.04) | 0.33(0.04) | 0.47(0.06) | 0.52(0.06) | 0.60(0.06) | 0.67(0.05) | – | 0.69(0.08) |

<sup>a</sup>Data presented as mean (standard deviation).
“–” indicates the corresponding IC was not found.

**Table 3.** The GoF, MWs and IIS of selected PMN/DMN under different combinations of model order and spatial smoothing levels using TCgICA.

| TCgICA |  | SM 0 |  | SM 6 |  | SM 9 |  | SM 12 |  |
| --- | --- | --- | --- | --- | --- | --- | --- | --- | --- |
|  |  | PMN | DMN | PMN | DMN | PMN | DMN | PMN | DMN |
| MO 20 | GoF | 0.39 | 0.49 | 0.42 | 0.43 | 0.43 | 0.42 | – | 0.46 |
|  | MW<sub>s</sub> | 7.30 | 6.54 | 7.73 | 6.74 | 7.71 | 6.93 | – | 7.53 |
|  | IIS<sup>a</sup> | 0.16(0.04) | 0.17(0.04) | 0.45(0.07) | 0.42(0.06) | 0.53(0.08) | 0.51(0.09) | – | 0.61(0.08) |
| MO 40 | GoF | 0.47 | 0.47 | – | 0.46 | 0.41 | 0.44 | 0.47 | 0.46 |
|  | MW<sub>s</sub> | 9.05 | 7.27 | – | 9.13 | 9.04 | 7.22 | 9.74 | 6.80 |
|  | IIS<sup>a</sup> | 0.08(0.03) | 0.09(0.03) | – | 0.27(0.06) | 0.40(0.07) | 0.40(0.09) | 0.48(0.08) | 0.45(0.09) |
| MO 60 | GoF | 0.47 | 0.55 | 0.49 | – | 0.48 | 0.35 | 0.46 | – |
|  | MW<sub>s</sub> | 10.56 | 7.60 | 11.15 | – | 11.39 | 7.45 | 11.05 | – |
|  | IIS<sup>a</sup> | 0.05(0.02) | 0.05(0.02) | 0.17(0.05) | – | 0.26(0.07) | 0.30(0.07) | 0.33(0.08) | – |
| MO 80 | GoF | 0.46 | 0.52 | 0.45 | 0.37 | 0.42 | – | 0.43 | – |
|  | MW<sub>s</sub> | 11.60 | 8.81 | 11.31 | 7.93 | 11.86 | – | 12.85 | – |
|  | IIS<sup>a</sup> | 0.02(0.01) | 0.02(0.01) | 0.05(0.03) | 0.05(0.03) | 0.09(0.06) | – | 0.16(0.09) | – |
| MO 100 | GoF | 0.46 | 0.51 | 0.41 | 0.44 | – | 0.33 | – | – |
|  | MW<sub>s</sub> | 11.76 | 9.12 | 10.65 | 7.94 | – | 9.88 | – | – |
|  | IIS<sup>a</sup> | 0.01(0.01) | 0.01(0.01) | 0.03(0.02) | 0.04(0.03) | – | 0.06(0.04) | – | – |

<sup>a</sup>Data presented as mean (standard deviation).
“–” indicates the corresponding IC was not found.

### Found or Not Found

As shown in Table 1, we did not identify PMN but successfully extracted DMN from the gRAICAR results using (low SM, low MO), including (0, 20) and (6, 20) combinations. Under all (low SM, high MO) combinations, both the PMN and DMN were successfully identified. In contrast, in the high SM conditions, PMN was not identified using the (9, 20) and (12, 20) combinations. DMN was not identified using the (9, 40) combination. The (12, 80) combination failed to identify DMN.

As shown in Table 2, we were unable to identify PMN from the IVA-GL results of the low MO conditions, such as (0, 20), (0, 40), and (6, 20). We did not find DMN in the (6, 40) condition. Under conditions of (low SM, high MO), we successfully identified both PMN and DMN. Under the high SM conditions, such as (9, 20), (12, 20), and (12, 40), IVA-GL failed to extract PMN from the rfMRI data. In the (high SM, high MO) conditions, the (12, 100) condition failed to identify PMN and the (9, 80) and (12, 80) conditions failed to identify DMN.

TCgICA failed to estimate the PMN component in the (6, 40) condition. TCgICA successfully detected both PMN and DMN using the (low SM, high MO) combination. In contrast, the high SM conditions, (12, 20), (9, 100), and (12, 100), and (9, 80), (12, 80), and (12, 100), failed to extract PMN and DMN, respectively.

In summary, only in the (low SM, high MO) condition, could both PMN and DMN be successfully separated from each other across all three algorithms. This finding indicates that PMN-DMN segregation is independent of the group ICA algorithm used. We also noticed that in the low MO condition, it was more difficult to identify PMN than DMN. However, as the spatial smoothing size increased, both PMN and DMN could not be identified due to the blurring effects of spatial smoothing. The spatial competition between PMN and DMN under conditions of high spatial smoothing became more evident when both PMN and DMN were successfully obtained. For instance, the posterior cluster size of DMN was greatly reduced due to the influence of PMN in the (9, 20), (9, 40) and (12, 40) conditions using TCgICA (shown in Supplementary Figure 3). This same effect was observed in the other two algorithms (gRAICAR and IVA-GL).

### GoF, MWs, and IIS

The (low SM, high MO) conditions always presented the highest GoF scores though the differences in GoF scores remained small across conditions. This may partly be due to the binarized templates and weighted RSN maps used in the analyses. MWs increased with model order, regardless of the algorithm used. Spatial smoothing can also raise the MWs, especially when the MOs are high.

Generally, IIS increased with spatial smoothing across the three ICA algorithms. The relationship between IIS and MO is algorithm-dependent. For gRAICAR and IVA-GL, IIS was not greatly influenced by MO, while for TCgICA, a higher MO produced lower IIS. For gRAICAR, the IIS differences between PMN and DMN in the high SM condition were relatively large, indicating that the two networks were not equally reproducible using this method. However, this was not the case for IVA-GL.

### Age effects

Given the age range of the 61 subjects are relatively wide, is it possible that the PMN-DMN segregation in the group-level is in fact introduced by the inter-individual variability? In order to tackle the problem, we re-did the analysis in the four extreme conditions, i.e. (SM 0, MO 20), (SM 0, MO 100), (SM 12, MO 20) and (SM 12, MO 100), on a sub-sample with 29 subjects and an age range of 20-30. The results obtained in the 61 subjects were replicated in this more homogenous sample. The results were presented in the Supplementary Figure 4.

### PMN-DMN overlaps at individual level

The group-level results showed significant spatial overlap between PMN and DMN, but it is unknown whether the PMN-DMN overlap presents at individual subjects or it is only due to averaging across subjects. This question is important as it helps to explain why the PMN-DMN segregation is rarely observed in seed-based functional connectivity studies. We therefore examined the individual-level variability of the PMN-DMN overlap in the condition of SM 0 and MO 100, which is the optimal combination to separate PMN and DMN according to the above results.

Individual-level ICs were first Z-transformed to have zero mean and unit standard deviation, and thresholded at Z > 2. We divided the number of voxels in the PMN-DMN overlap by the number of voxels in PMN, yielding an overlap percentage for each subject. The mean and standard deviation of this index in gRACAR, IVA-GL, and TCgICA were 12.1 (4.4), 17.7 (3.4) and 8.2 (5.6) respectively. The fact that the standard deviation is small relative to the mean of the overlap percentage indicates that the PMN-DMN overlap can be stably observed across subjects.

## Discussion

We examined the effects of spatial smoothing and model order on segregation between PMN and DMN using group ICA. We also took the type of algorithm used into consideration. Our findings demonstrated that these two networks can be reliably functionally segregated using a combination of low-level spatial smoothing during preprocessing and high model order in the ICA.

Looking through the DMN-related studies, it is not difficult to find an RSN with a spatial configuration similar to PMN that was identified as (posterior) DMN [31, 32]. In addition, some studies attributed the precuneus/posterior cingulate cortex group differences to DMN [33]. The distinction between PMN and DMN is important for determining the role of DMN, as DMN is related to many brain disorders [34, 35]. For instance, disruptions in DMN have been consistently reported in AD (Alzheimer’s disease) [31, 36, 37]. PMN is related to the processing of novel memories. A core symptom of AD is memory loss; therefore, it is reasonable to inquire whether PMN is also involved in the functional pathology of AD.

Both PMN and DMN could only be identified when a combination of low spatial smoothing and high model order were used. Under these conditions PMN and DMN had the highest GoF across all three algorithms. Previous studies have shown that more fine-grained RSNs can be extracted when using a high model order [10]. This was replicated in the present study. In fact, as the model order increased, many more RSNs emerged, the majority of which are still functionally unfamiliar to the community.

Spatial smoothing has been found to reduce functional specificity, for instance, merging two activation clusters into one [13], though its impact on group ICA is rarely evaluated. As spatial smoothing size became larger, PMN and DMN could no longer be successfully separated. Our findings indicate that one should be careful when using spatial smoothing, especially when functional specificity is a priority. In the past, low model order and middle-to-high spatial smoothing were widely applied in group ICA analysis [11, 38]. Under these conditions PMN and DMN can hardly be separated simultaneously, complicating the interpretation of these findings.

To rule out the possibility that the PMN-DMN segregation was totally introduced by the inter-individual variability of an RSN, for instance, DMN may have large variability in different ages, we replicated the analysis in a more homogenous sample. Our results and the conclusion still hold (see Supplementary Figure 4): PMN and DMN can be stably separated only in the condition of low spatial smoothing and high model order.

Currently, there are different ways of performing group-level ICA applying different assumptions to the data. To mitigate algorithm-dependent effects, three typical group ICA algorithms were evaluated in the present study. We observed that the three algorithms converged upon the same conclusion regarding PMN and DMN segregation, although there were algorithm-dependent effects on inter-individual reproducibility. Using both simulated and real data, previous studies have compared IVA-GL and TCgICA, concluding that IVA-GL outperformed TCgICA in capturing inter-individual variability [39, 40]. We also found relatively higher inter-individual similarity using IVA-GL when compared to the other two algorithms. However, due to the lack of a ground truth, the difference in inter-individual similarity across algorithms should be interpreted cautiously. Therefore, we recommend the use of a combination of multiple group ICA algorithms or group ICA with other non-ICA algorithms to increase the robustness and reliability of the findings, a major concern in neuroimaging.

Individual variability of resting-state networks is of great interest and the IIS was indeed observed to vary with the choice of MO and SM. IIS increased with SM independent of algorithms. Spatial smoothing makes the RSNs “larger” (more voxels passed the threshold), and thus more common voxels will be obtained among individuals, which would result in higher IIS. IIS decreased with MO in TCgICA, while kept relatively stable with MO in gRAICAR and IVA-GL. It is possible that TCgICA makes a higher homogeneity assumption among individuals than the other two algorithms. In the low MO, each RSN incorporates many brain regions, which will be split into functionally more homogenous smaller RSNs with the increase of MO. As a result, in the higher MO, the homogeneity assumption in TCgICA may not hold, leading to lower IIS.

While seed-based functional connectivity has been widely employed by the neuroimaging community due to its ease of both understanding and performance, many studies demonstrated its high dependencies on strategies of seed-selection, seed-location, and seed-shape. Spatial overlaps between PMN and DMN at the individual level were stably observed across different algorithms. Thus, it is of challenge to apply seed-based functional connectivity analysis method to investigating PMN-DMN difference, due to the ambiguity in defining seed regions. Patten analysis such as ICA provides an effective approach to segregate the two networks. Here, we demonstrate a combination of group ICA parameters to achieve optimal segregation of PMN and DMN.

In this study, we did not perform group ICA multiple times to obtain robust RSNs as in some previous studies [9, 10]. The considerations behind this were two-fold. First, performing multiple runs of ICA and combining these results is a challenging task in that of itself and will induce unexpected biases while offering only limited improvements [41-43]. Second, group-level ICA algorithms pool all individual datasets and can efficiently counterbalance the inherent indeterminacy of ICA analysis.

As both PMN and DMN include precuneus and posterior cingulate cortex, it is less likely to set them apart based on anatomical labels. The most obvious feature and difference between PMN and the posterior part of DMN lie in the spatial configuration. DMN has one cluster, which was flanked by two clusters in PMN in the mid-line slice. The aim to set the anchor points is to characterize this kind of spatial features. The only uncertainty probably caused by the anchor points is in the situation where PMN or/and DMN were labeled as “Not-Found”. To confirm that the “Not-Found”-labeled PMN and DMN were truly non-existent, we selected PMN and DMN only by the highest correlation with the templates and visually validated all the results. The results were presented in the supplementary materials.

In practice, other factors are also worth considering, for instance, data quality and hypothesis testing methods. The common use of spatial smoothing is based upon the Gaussian kernel, which can blur the boundaries between different signals. Recent advances in non-local smoothing may improve PMN-DMN segregation due to the high level of spatial smoothing [44, 45]. High model order and low spatial smoothing may not be noise-resistant enough and can therefore be sub-optimal if the fMRI data has a limited number of time points or low signal-to-noise ratio.

In the current study, the subjects were scanned with an eye-closed resting-state condition and it is worthwhile to examine the effects among different resting-state conditions. However, we believed that the current results would not likely to be greatly influenced by different resting conditions, i.e. eye-open versus eye-closed. Firstly, the PMN-DMN segregation has been identified in some studies where eye-open resting state fMRI datasets were utilized [5, 7, 8]; Secondly, previous studies showed that minor changes took place in resting-state networks among different conditions [46].

Taken together, our study demonstrated that low spatial smoothing and high model order is optimal when using group ICA to segregate PMN from DMN. Under these conditions, both PMN and DMN can be detected and extracted without interference. Goodness-of-fit were highest in these conditions when compared to the other conditions. Inter-individual reproducibility varied across algorithms. Data quality and smoothing methods must be considered in setting parameters to ensure the validity and robustness of any derived findings.

## Acknowledgements

This work was supported by the National Basic Research (973) Program (grantnumber:2015CB351702), the Natural Science Foundation of China (grant numbers: 81571756, 81270023, 81278412, 81171409, 81000583, 81471740, 81220108014), Beijing Nova Program (XXJH2015B079 to Z.Y.), the Outstanding Young Investigator Award of Institute of Psychology, Chinese Academy of Sciences (to Z.Y.), the Key Research Program and the Hundred Talents Program of the Chinese Academy of Sciences (KSZD-EW-TZ-002 to X.N.Z).

